# Accelerating scRNA-seq Analysis: Automated cell type annotation using representation learning and vector search

**DOI:** 10.1101/2025.10.06.680787

**Authors:** Stephen R. Williams, Fedor Grab, Govinda M. Kamath, Yerdos Ordabayev, Jeff Mellen, Patrick Roelli, Kristian Cibulskis, Erik Lehnert, Fen Xie, Miguel Covarrubias, Nur-Taz Rahman, Timothy Tickle, Emre Erhan, Nicolas Malfroy-Camine, Kevin Lydon, Mehrtash Babadi, Nigel F. Delaney

## Abstract

Cell type annotation in single-cell RNA sequencing (scRNA-seq) experiments is the fundamental step of assigning cell types to individual cells or clusters of cells based on their gene expression profiles. This process is crucial for developing biological insights from scRNA-seq experiments. We present a service that automates cell type annotation for 10x Genomics single-cell gene expression samples, enabling researchers to rapidly and accurately categorize cells within a sample. This service operates on the basis of reverse search: it compares each cell’s gene expression profile against the Chan Zuckerberg CELL by GENE (CZ CELLxGENE) Census, a comprehensive repository of published scRNA-seq datasets enriched with community-annotated cell types, and yields cell type annotations through summarizing the labels associated with similar cells. The annotation algorithm employed in this service avoids reliance on predefined marker genes or tissue-specific references, providing both fine-grained and coarse annotations. These initial annotations can be further refined by investigators to suit their specific research needs.

## 1 Introduction

Since the advent of bulk RNA-sequencing [1], a critical initial step in analyzing gene expression data has been to determine the composition of cell types within a sample. Bulk RNA-sequencing studies relied on deconvolution methods to infer the cellular composition of the tissue of interest [2, 3]. The emergence of single-cell RNA sequencing (scRNA-seq) technologies has revolutionized this field, enabling the characterization of cellular heterogeneity with unprecedented resolution and scale. Despite these advancements, accurately annotating individual cells within scRNA-seq datasets remains a significant analytical challenge, often requiring substantial time and effort.

In an effort to characterize the tremendous diversity of cell types in the human body, large-scale collaborative efforts have emerged to generate comprehensive reference datasets. Initiatives like the Human Cell Atlas [4] and Tabula Sapiens [5] aim to create a comprehensive map of all human cell types by generating and integrating scRNA-seq data from diverse tissues and individuals. By building upon these extensive reference datasets, researchers can develop and leverage powerful computational methods and machine learning algorithms to accurately and efficiently annotate cells within their own experiments, accelerating the pace of scientific discovery in the field of single-cell genomics.

The Chan Zuckerberg Initiative Cell by Gene (CZ CELLxGENE) program [6, 7] aims to unify and expand upon these efforts by creating a centralized resource of scRNA-seq data. This initiative integrates data from various sources, including the Human Cell Atlas, Tabula Sapiens, and individual contributors to build a comprehensive and accessible resource of annotated cell types. Annotations from diverse datasets are easily leveraged because they use shared ontology framework, the Cell Ontology. [8]. The CZ CELLxGENE program not only provides a valuable resource for researchers but also serves as an ideal foundation for developing and training machine learning models for cell type annotation. By utilizing this rich and diverse dataset, researchers can train algorithms that can accurately predict cell types based on gene expression patterns, ultimately accelerating the pace of scientific discovery in single-cell genomics.

Current cell annotation methods often present significant challenges [9, 10]. Many existing methods require specialized bioinformatics expertise to implement and interpret, hindering their accessibility to researchers from non-technical backgrounds. Additionally, many methods are developed for specific tissue types, limiting their generalizability. Furthermore, some approaches rely heavily on predefined marker genes, which may not always accurately capture the full spectrum of diversity within a specific cell type. Finally, label transfer methods, while promising, can be susceptible to inaccuracies if the reference dataset is not sufficiently similar to the target dataset, potentially leading to erroneous cell type assignments.

Here, we present a machine learning reverse search methodology that leverages the entirety of the 10x Genomics data available in CZ CELLxGENE Census for accurate and automated cell type annotation. This reference dataset encompasses a vast and diverse collection of human and mouse single-cell RNA-sequencing data. By benchmarking our annotation service on independently held-out datasets, we demonstrate high accuracy in annotating cells across diverse tissue types and experimental modalities, including both single-cell and single-nucleus assays. Furthermore, we have integrated our proposed framework into services that are readily accessible to researchers without the need for extensive computational resources or complex workflows.

## 2 Methods and Results

The proposed automated cell annotation framework consists of the following stages (Fig 1): (1) An appropriate low-dimensional embedding model is trained on the entire reference data collection to obtain compact vector representations of each reference cell; (2) The vector representations are used to build a fast approximate nearest-neighbor (ANN) lookup index; (3) At inference time, the gene expression profile of each of the user-provided cells is transformed into the same embedding space and matched to transcriptionally similar reference cells via ANN; (4) Finally, the labels of similar reference cells are aggregated and post-processed into a cell type label proposal. One’s choice of reference data pre-processing and embedding (i.e., feature selection, transformation, embedding dimensionality) is expected to have significant ramifications for the overall accuracy performance of the framework. In the next few subsections, we detail such modeling choices.

**Fig. 1:**
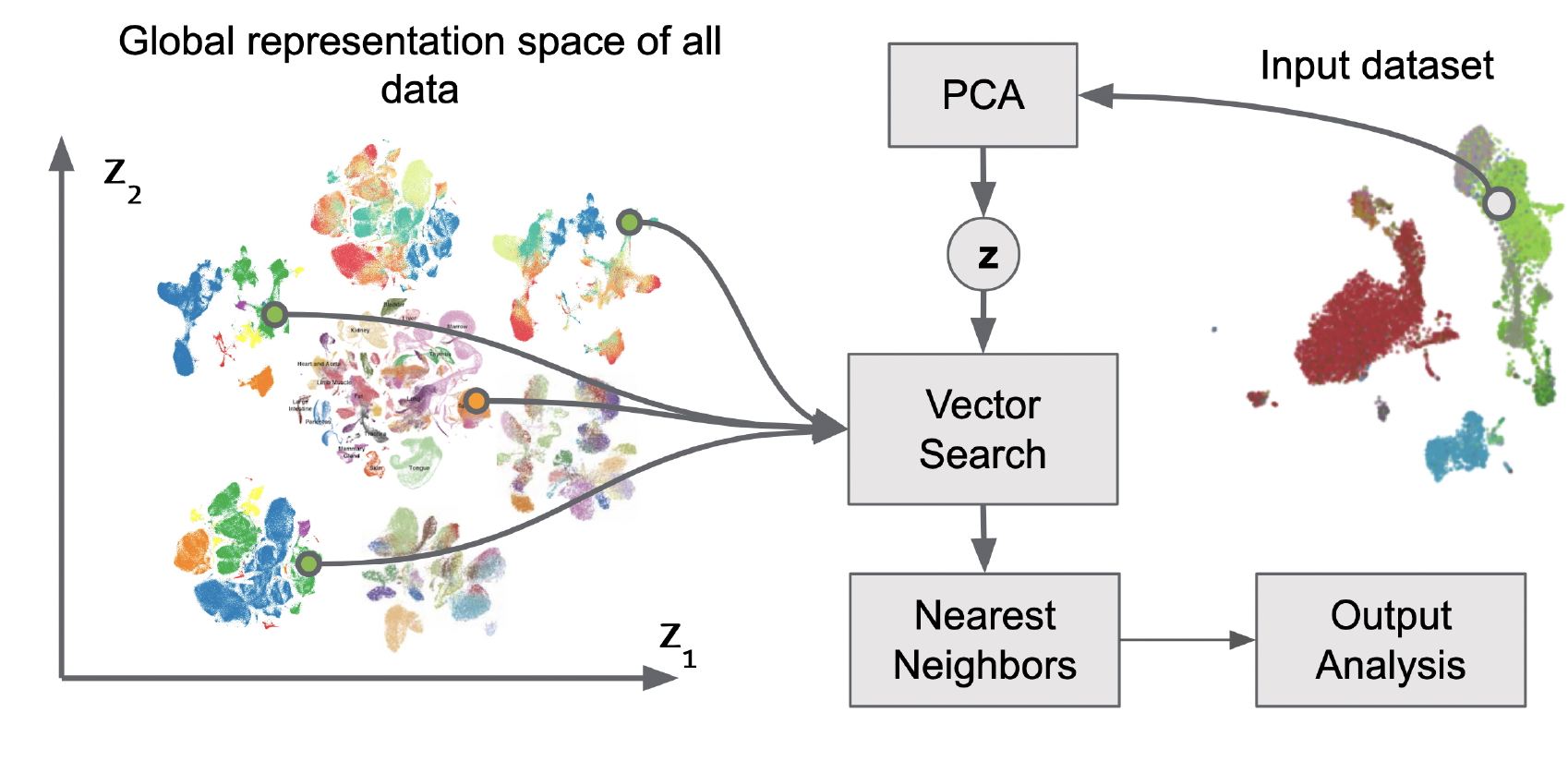
Workflow for the proposed referenced-based cell annotation via reverse search. Input data undergoes dimensionality reduction via Principal Component Analysis (PCA), generating a lower-dimensional vector embedding *z*. ANN vector search within this global representation space identifies nearest neighbors to the query cell, which are used to assign cell annotations.

### 2.0.1 Feature Selection

For each species in question (human and mouse), we start with Ensembl gene IDs filtered according to Cell Ranger recommendations. These correspond to GRCh38-2020-A (Human) and mm10-2020-A (Mouse) transcriptome references. We consider various subsets of these initial gene sets as input to downstream modeling as follows:

- *Full Schema*: Includes all genes from the GRCh38-2020-A (Human) and mm10-2020-A (Mouse) reference transcriptome annotations without further filtering.
- *Top 10,000 Highly Variable Genes (HVG)*: Subsets the reference gene set to the top 10,000 HVG based on normalized dispersion computed over the entire CZ CELLxGENE Census data catalog. The HVG selection strategy is identical to the approach introduced by the R package Seurat [11].
- *Highly Variable Genes with mean and normalized dispersion cutoffs:* Selects HVG based on cutoffs imposed over log-normalized mean gene expression and quantile-normalized dispersion computed over the entire CZ CELLxGENE Census data catalog. We specifically use the default values set in scanpy[12] for the cutoffs: minimum mean log expression: 0.0125, maximum mean log expression: 3.0, minimum normalized dispersion: 0.05.

### 2.0.2 scRNA-seq Data Preprocessing Transformations

We perform standard library size normalization on all model variants. This operation is defined as:

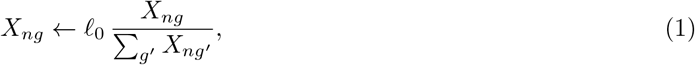

where *ℓ*_0_ = 10, 000 is the target library size. Crucially, this operation is performed before feature selection, such that the total count (denominator) is calculated over all measured genes. We then (optionally) perform a series of additional transformations. These include: (1) logarithmizing the counts (“log1p”):

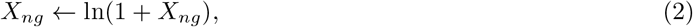

where we add a pseudo-count of 1 in order to keep genes with 0 expression at 0 after transformation; (2) column-wise *z*-scoring:

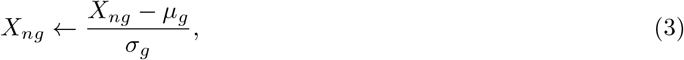

where *µ*_*g*_ and *σ*_*g*_ denote the mean and standard deviation, respectively, calculated over all cells in the data catalog; (3) Scaling by median of non-zero counts:

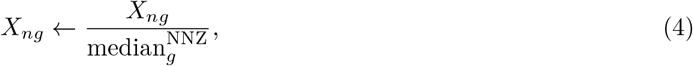

where 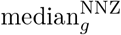 denote the median of non-zero counts of gene *g* calculated over all cells in the data catalog. We consider mixing and matching these transformations aimed to identify the most effective approach for preparing the data for dimensionality reduction. In particular, we consider the following preprocessing transformation combinations: (1) No additional preprocessing beside library size normalization; (2) Only log1p transformation; (3) Only *z*-score transformation; (4) log1p and *z*-score; (5) Scaling by median of non-zero counts; (6) log1p and then scaling by median of non-zero log1p counts.

### 2.0.3 Embedding using Principal Component Analysis (PCA)

We employed Principal Component Analysis (PCA) as the embedding model, which is a commonly-used and highly-effective strategy for scRNA-seq dimensionality reduction. To enable large-scale data processing, we implemented a distributed and incremental implementation of PCA [13, 14] as a part of the open-source Cellarium ML library [15]. We used this scalable implementation to separately obtain PCA embeddings for human and mouse data. In total, we generated 72 different model variations and benchmarked each variation separately for each organism. These combinatorial variations were defined by 4 choices of PCA embedding dimensionalities (64, 128, 256, 512), 3 choices for feature selection, and 6 choices for data preprocessing transforms detailed above. Our internal benchmarks (not reported here) identified the most effective strategy separately for human and mouse cell type annotation. More concretely, our human scRNA-seq embedding model uses all genes (full schema), top 512 principal components, log1p transformation, and *z*-score normalization. Our mouse scRNA-seq embedding model uses top 10,000 HVGs, log1p transformation, and *z*-score normalization. We will present benchmarking results obtained using these embeddings in the next sections.

### 2.0.4 Approximate Nearest Neighbor (ANN) Search

To assign fine-grained cell type annotations to cells, the service uses an ANN lookup strategy. This approach efficiently searches the space of gene expression profiles to identify the closest matching reference cells using the aforementioned low-dimensional embeddings as a proxy for gene expression similarity. By comparing the gene expression profile of a query cell to the profiles of known cell types in the reference dataset, the model can assign a fine-grained cell type annotation based on the nearest neighbor’s identity. We use ScaNN [16–18] to query and return the top 500 approximate nearest neighbors from the model and summarize these data, including the number of cells assigned to the reference cell type and the CZ CELLxGENE Census dataset(s) from which that assignment came.

The key advantage of using ANN over K-Nearest Neighbors (KNN) lies in computational efficiency, especially crucial when dealing with large-scale scRNA-seq datasets comprising 100M+ cells. While KNN provides the most accurate results in principle, ANN methods combined with dimensionality-reduced representations offer a crucial trade-off: they sacrifice absolute accuracy for a significant gain in computational speed. In the context of large-scale scRNA-seq datasets, this speedup is critical for enabling rapid and efficient cell type annotation. Furthermore, given the inherent noise and dropout in gene expression profiling, maintaining absolute accuracy in identifying the closest gene expression matches is unlikely to yield a significant advantage.

### 2.0.5 Tree Representation of Cell Type Ontological Relationships

Commonly used cell types terms have different degrees of granularity (e.g. monocyte vs. non-classical monocyte) and are ontologically related to one another. The ontological relationships between cell type terms (e.g. non-classical monocyte is_a monocyte) are captured in multiple community-driven efforts, most prominently, the cell type ontology project (CL) [19] which is the ontology adopted by CZ CELLxGENE Census. We leverage this resource to aggregate cell type terms obtained through ANN lookup into more canonical cell type terms. In order to facilitate interpretation and readability of the results, we generate a subtree of the full CL graph. First, we subset CL by all cell types that are present in our single-cell reference dataset. We then manually pick a set of ontology terms that are representative of the major cell types that are commonly used and well understood. These cell types are defined as “coarse”, also referred to as “display nodes”, that form the base tree. Finally, all cell types were aggregated to one unique parent display node present in the base tree. If a cell type has multiple parents in the original CL graph, a representative parent was manually chosen. This process results in a tree with all cell types present in the model and a mapping from all fine-grained cells to the corresponding coarse cell type. The tree is provided at https://10xgen.com/cell-ontology-tree.

Following the ANN lookup and assignment of fine-grained cell type annotations, the results are further summarized by mapping each fine-grained cell type to a corresponding coarse display node. The tree allows for a more concise and interpretable overview of the cell type assignment (*e*.*g*., ‘T-cell’ instead of ‘CD8-positive T-cell’ and ‘B cell’ instead of ‘Naive B cell’), enabling researchers to easily navigate their results while at the same time having the primary fine-grained annotation at their disposal. This hierarchical representation facilitates a clear understanding of mappings for each cell.

### 2.1 Benchmarking

To assess model performance, we benchmarked our proposed annotation framework across various experimental parameters. We trained separate models (as described earlier) on the same CZ CELLxGENE Census version “2024-05-06”, excluding a number of datasets from specific organ systems (blood, brain, heart, kidney, lung), assay versions, and suspension types (cells and nuclei). This approach ensured that model performance was evaluated on independent datasets, minimizing potential biases.

For benchmarking, 26 human and 5 mouse datasets were selected representing a combination of assays, tissues, and suspension types (see Table 1 in Appendix). Models for benchmarking were trained on the reference datasets, excluding benchmarking datasets, non-10x assays, and non-primary data from the CZ CELLxGENE Census (Tables 1 & 2).

**Table 1:** Summary statistics of datasets used to train the human cell type annotation model.

| Assay | Suspension Type | # Cells |
| --- | --- | --- |
| 10x 3' Transcription Profiling | Cell | 165,055 |
| 10x 3' v1 | Cell | 54,589 |
| 10x 3' v2 | Cell | 6,621,509 |
| 10x 3' v3 | Cell | 5,471,480 |
| 10x 5' Transcription Profiling | Cell | 230,657 |
| 10x 5' v1 | Cell | 3,709,801 |
| 10x 5' v2 | Cell | 1,962,206 |
| 10x 3' v2 | Nuclei | 631,784 |
| 10x 3' v3 | Nuclei | 12,138,706 |
| Total |  | 30,985,787 |

**Table 1:**
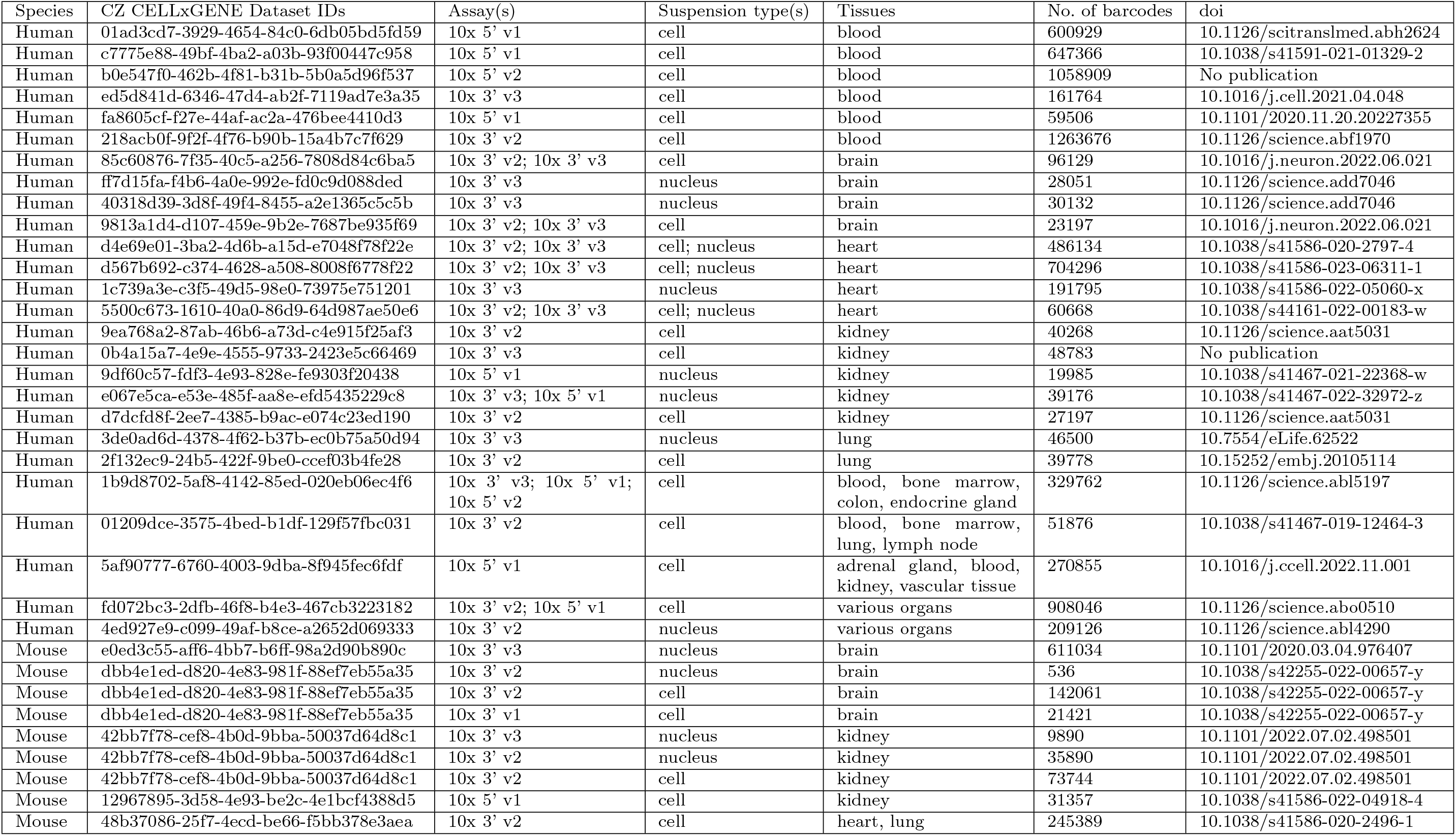
Datasets used for benchmarking.

**Table 2:**
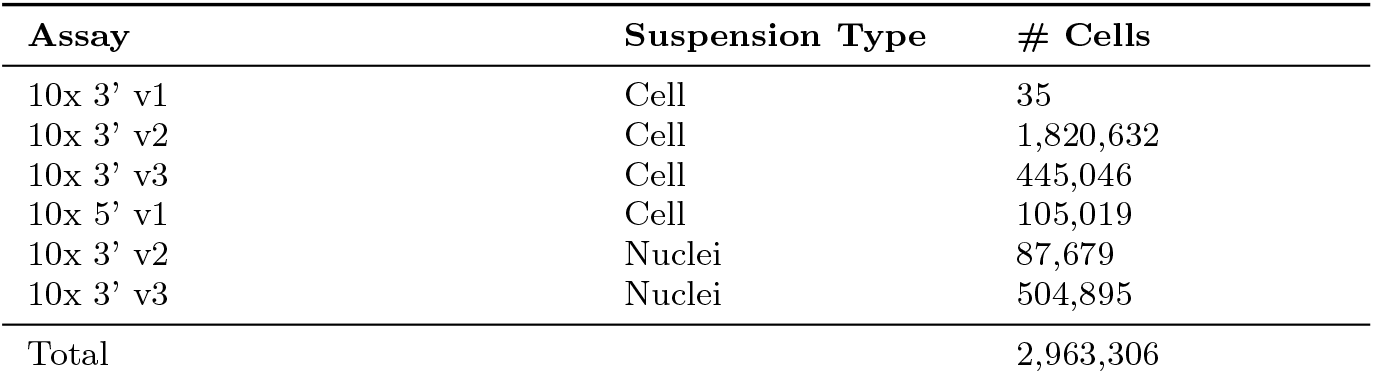
Summary statistics of datasets used to train the mouse cell type annotation model.

To estimate the error rate in the models, the benchmarking datasets were annotated using the mouse and human benchmarking models. A cell was considered correctly annotated if its coarse cell type was consistent with the community-provided annotation mapped to its coarse annotation. For example, ‘Activated CD4-negative, CD8-negative Type I NK T cell’ and ‘Activated CD4-positive, alpha-beta T cell’ both map to ‘T cell’ and would be considered concordant. The decision to not judge concordance on the raw result of the benchmarking models was due to the uneven granularity of annotation used in different reference datasets.

#### 2.1.1 Benchmarking Performance by Assay Version

Table 3 and Table 4 present benchmarking results by assay version for human and mouse data, respectively (Fig. 2a). Overall, 10x 3’ v3 demonstrated the highest mean concordance rate (96.7%) for both human and mouse annotation models, suggesting superior performance compared to other assay versions.

**Table 3:** Benchmarking datasets stratified by assay version (human)

| Assay Version | # Datasets | Mean Concordance rate |
| --- | --- | --- |
| 10x 3' v2 | 6 | 93.2% |
| 10x 3' v2 10x 3' v3 | 5 | 94.2% |
| 10x 3' v2 10x 5' v1 | 1 | 86.8% |
| 10x 3' v3 | 6 | 96.7% |
| 10x 3' v3 10x 5' v1 | 1 | 96.7% |
| 10x 3' v3 10x 5' v1 10x 5' v2 | 1 | 98.2% |
| 10x 5' v1 | 5 | 92.7% |
| 10x 5' v2 | 1 | 99.2% |

**Table 4:**
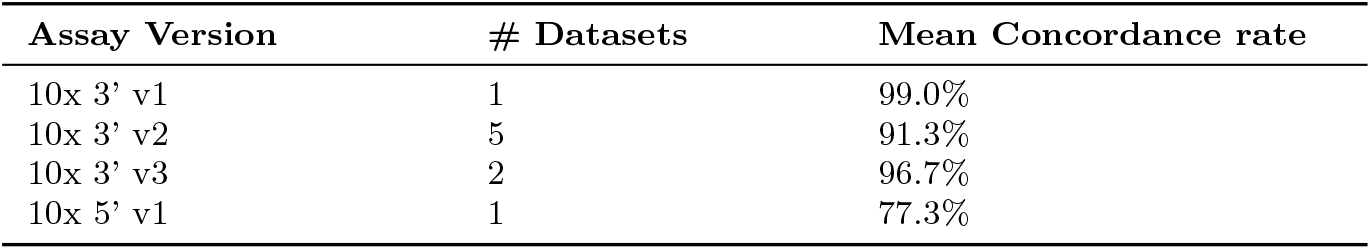
Benchmarking datasets stratified by assay version (mouse)

**Fig. 2:**
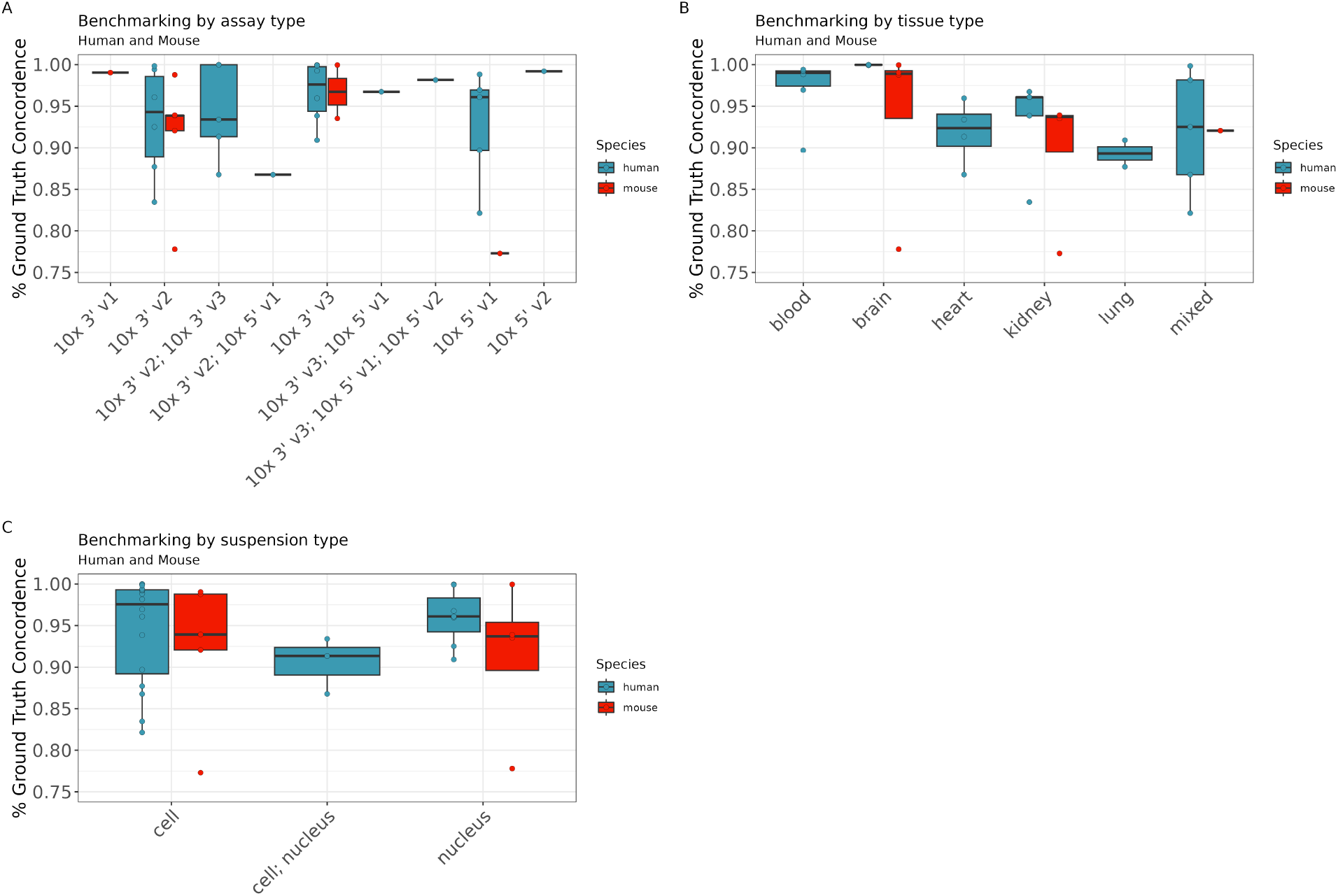
Benchmarking: Human and Mouse. Human shown in red mouse shown in blue. Each point represents a single study which was held out during model development and used for benchmarking. (A) Concordance of cell annotation results as compared to ground truth by 10x Genomics assay as defined by CZ CELLx-GENE Census. Assays names separated by a semicolon represent studies that combined multiple 10x assays. (B) Concordance of cell annotation results as compared to ground truth by tissue type as defined by CELLx-GENE. Mixed represents studies which comprise more than one of the the benchmarking tissue types (blood, brain, heart, kidney, or lung. (C) Concordance of cell annotation results as compared to ground truth by suspension type as defined by CELLxGENE. Suspension types separated by a semicolon represent studies which have mixtures of suspension types (cell and nucleus).

#### 2.1.2 Benchmarking Performance by Tissue Type

Table 5 and 6 show benchmarking results by tissue type (Fig. 2b)). While performance was high across most tissues, brain tissue consistently exhibited high concordance rates in both human and mouse datasets. In contrast, lung tissue showed lower concordance in the human dataset. This may be due to the presence of cell types that are challenging to annotate, or may reflect the effect of which display nodes were selected for each cell type (in general, more specific display nodes tend to have lower concordance rates).

**Table 5:** Benchmarking datasets stratified by tissue type (human)

| Tissue Type | # Datasets | Mean Concordance rate |
| --- | --- | --- |
| blood | 6 | 97.2% |
| brain | 4 | 100.0% |
| heart | 4 | 91.9% |
| kidney | 5 | 93.2% |
| lung | 2 | 89.3% |
| mixed | 5 | 91.9% |

**Table 6:** Benchmarking datasets stratified by tissue type (mouse)

| Tissue Type | # Datasets | Mean Concordance rate |
| --- | --- | --- |
| brain | 4 | 93.9% |
| kidney | 4 | 89.7% |
| heart & lung | 1 | 92.1% |

#### 2.1.3 Benchmarking Performance by Suspension Type

Tables 7 and Table 8 present benchmarking results by suspension type (cells and nuclei, Fig. 2c). In general, performance was comparable between cell and nuclei suspensions, with slight variations observed across species.

**Table 7:** Benchmarking datasets stratified by suspension type (human)

| Suspension Type | # Datasets | Mean Concordance |
| --- | --- | --- |
| Cell | 16 | 94.5% |
| Nucleus | 7 | 96.0% |

**Table 8:** Benchmarking datasets stratified by suspension type (mouse)

| Suspension Type | # Datasets | Mean Concordance |
| --- | --- | --- |
| Cell | 5 | 92.2% |
| Nucleus | 4 | 91.3% |

Overall, the results demonstrate high accuracy and generalizability of the developed annotation models across different experimental conditions, including variations in assay versions, tissue types, and suspension types. These findings highlight the robustness and potential broad applicability of the models in diverse scRNA-seq experiments.

#### 2.1.4 10x Genomics Flex Gene Expression

To further assess the generalizability of our models, we evaluated their performance on 10x Genomics Flex probe-based whole genome single-cell assays [20], which were not present in the training data. To assess performance on Flex gene expression samples, the results of cell annotation were manually compared to cell type annotations provided by Azimuth [21]. The mean concordance across lung, kidney, breast, pancreas, brain, lymph node, and liver samples tissues was 76%. The lower performance may partially have been due to challenges consistently mapping cell types across the two datasets. The models demonstrated reasonably strong performance on these assay types, indicating their robustness and adaptability beyond the specific datasets used for training. This finding suggests that the developed models can be effectively applied to a broader range of single-cell RNA-sequencing technologies with appropriate considerations of caveats (see Discussion).

### 2.2 Model Updates after Benchmarking

We observed that the mouse benchmarking datasets comprised nearly one-third of all mouse data in the CZ CELLxGENE Census, whereas the human benchmarking datasets formed a negligible fraction of the total human data. This imbalance arises because the Census contains substantially more human cells than mouse cells. Given this disparity, we determined that reintegrating the mouse benchmarking datasets would significantly boost our final system’s performance.

Following our benchmarking experiments and in preparation for the release, we retrained the mouse embedding models and rebuilt the ANN search indices using the entire CZ CELLxGENE Census mouse dataset, including those previously reserved for benchmarking. A manual review of multiple internal datasets confirmed that the updated mouse annotation model either maintained or improved annotation quality, especially for tissue types that benefited the most from the expanded training set. This new mouse model is now deployed in our user-facing software.

Because the human benchmarking datasets accounted for only a minor share of the total human reference data, we decided not to incorporate them into the current release. However, we plan to include them in a future update, where they will be used to retrain and enhance the human model.

### 2.3 Running Cell Annotation

Both the Broad Institute and 10x Genomics provide users an ability to run cell annotation, and there are 4 ways to run cell annotation using the methods described here. Non-interactively using the 10x Genomics Cell Ranger software and via the count, multi, and annotate pipelines (Cell Ranger 9.0+) and interactively within Jupyter notebooks or standalone Python scripts using the Cellarium CAS [22].

The Cell Ranger pipeline offers both local and cloud-based execution options. For local execution, the process involves three core pipelines: cellranger count, cellranger multi, and cellranger annotate. Alternatively, the pipeline can be executed on the 10x Genomics Cloud Platform, providing scalability and eliminating the need for local computational resources or the use of a CLI. Cloud-based execution requires setting up cloud authentication as outlined in the Cell Ranger documentation.

When running cell annotation using cellranger count, multi, or annotate pipelines locally the stage which performs cell annotation makes a user-authorized call out to the 10x Genomics Cloud Platform where the feature/count matrix (HDF5 format[23]) is uploaded and cells are labeled using the process shown in Fig. 1. When running cell annotation in the 10x Genomics Cloud Platform the user has already authorized their data to be stored by 10x Genomics and cell annotation is run automatically.

When running cell annotation using Cellarium CAS while following the user guide [22], users must obtain a token from the Cellarium AI team by accessing the Cellarium CAS public beta signup form.

## 3 Discussion

Cell type annotation remains one of the first and most challenging steps of single-cell gene expression analysis. There is both a huge breadth of cell types as well as different means of classifying them (e.g., cellular state, morphology, connectivity to other cells), which makes a one-size-fits-all solution challenging. By providing cell annotation through Cell Ranger and Cellarium CAS, the goal is for researchers to be able to start investigating their dataset and refining annotations without the need to curate lists of marker genes, identify tissue-specific references, or develop scripts to perform annotation. However, this approach does require refinement on the part of researchers and the following caveats and biases should be kept in mind.

### 3.0.1 Caveats & Biases

While benchmarking demonstrated that the models performed well in a range of assay and tissue types, biases are present that can impact the accuracy of cell type calls. First and foremost, accuracy is limited by the presence of the appropriate cell types in the CZ CELLxGENE Census reference. For example, cell lines are not included, and organoids and diseased tissues are relatively poorly represented compared to healthy tissue. Biases in the composition of the census may be improved by periodic releases of the model with new datasets that include these under-represented sample types. However, we recommend that researchers with uncommon cell types or states that are expected to have large impacts on transcript prevalence (e.g., cancer) check that the version of CZ CELLxGENE used to build the model contains representative datasets.

A second source of bias is the lack of uniform levels of annotation and relative dataset sizes. For example, large datasets with less refined annotations present in the reference can result in less cells annotated with less specific cell types. Future improvements to investigate may include preferential inclusion of datasets with more specific cell annotations or sub-sampling to mitigate the effect of relative dataset sizes.

Finally, although efforts have been made to produce a robust model, misannotation does occur at a low rate. We have noted that this tends to be pronounced in low-quality samples or cells with low UMI counts, but some occurrences may be due to misannotated cells in the reference. We recommend verifying any unusual or rare cell types using marker genes when available.

### 3.0.2 Future Improvements

Cell annotation remains an active area of research with numerous methods available and active research into new approaches. A key rationale for developing a cloud-based approach to annotation is the ability to add new or updated versions of existing models without requiring updates to the Cell Ranger pipeline. We encourage researchers who benefit from a robust starting point for cell annotations to, when permissible, refine annotations and contribute datasets to public repositories to improve future models.

## 4 Acknowledgments

We thank the Chan Zuckerberg Initiative CELLxGENE team, in particular Ambrose Carr, for their advice and support for this project.

## 7 Supplementary data

For a comprehensive view of the cell ontology tree, we refer readers to the detailed hierarchical structure available at https://10xgen.com/cell-ontology-tree.

